# Community Engagement during outbreak response: standards, approaches, and lessons from the 2014-2016 Ebola outbreak in Sierra Leone

**DOI:** 10.1101/661959

**Authors:** Jamie Bedson, Mohamed F. Jalloh, Danielle Pedi, Saiku M. Bah, Katharine Owen, Allan Oniba, Musa Sangarie, James Fofanah, Mohamed B. Jalloh, Paul Sengeh, Laura A. Skrip, Benjamin M Althouse, Laurent Hébert-Dufresne

**Author notes:** Corresponding author: Benjamin M Althouse, Institute for Disease Modeling, 3150 139th Ave SE, Bellevue, WA, 98005. These authors contributed equally (Jamie Bedson, Mohamed F. Jalloh). Co-senior authors (Benjamin M Althouse, Laurent Hébert-Dufresne).

## Abstract

- The Social Mobilization Action Consortium (SMAC) was Sierra Leone’s largest coordinated community engagement initiative during the 2014 - 2016 Ebola outbreak. It worked in all 14 districts in Sierra Leone across >12,000 communities (approximately 70% of all communities), through 2,466 trained Community Mobilizers, a network of 2,000 mosques and churches, and 42 local radio stations.
- We describe SMAC’s Theory of Change and utilization of the Community-Led Ebola Action (CLEA) approach. We present an extensive dataset of community engagement and monitoring with a focus on over 50,000 SMAC weekly reports collected by Community Mobilizers between December 2014 and September 2015.
- Community engagement and real-time data collection at scale is achievable in the context of a health emergency if adequately structured, managed, coordinated and resourced.
- We describe a correlation between systemic community engagement, community action planning and Ebola-safe behaviors at community-level.
- The SMAC integrated approach demonstrates the scope of data – including surveillance data - that can be generated directly by communities through structured community engagement interventions implemented at scale during an Ebola outbreak.
- We highlight important insights gleaned over time on how to informally integrate social mobilization into community-based surveillance of sick people and deaths.

## The Need for Integrated Data-Driven Community Engagement in Health Emergencies

Community engagement during public health emergencies is increasingly recognized as an important component to foster enabling and reinforcing conditions for behavior change to reduce the spread of disease [1, 2]. In 2009, the World Health Organization (WHO) convened an informal consultation to develop standards and identify best practices for social mobilization in public health emergencies [1]. The consultation concluded that there was a general under-appreciation of the behavioural imperative that underlies responses to public health emergencies, despite the fact that human behaviour drives epidemic emergence, transmission and amplification. An interagency guide on communication for behavioural impact (COMBI) during outbreak response was then shortly developed by WHO, UNICEF, and partners in 2012. Since then, the recognition of the critical role of community engagement in disease response has been reflected in a range of international agreements and guidelines [3, 4].

The importance of community engagement was uniquely exemplified during the 2014-2016 outbreak of Ebola Virus Disease (Ebola) in West Africa, which resulted in nearly 30,000 cases in Guinea, Sierra Leone, and Liberia [5]. Sierra Leone was hit the hardest with 14,124 cases and 3,956 deaths attributed to Ebola [6]. As numbers of cases rapidly increased there was a growing consensus that behavior change was required to reduce complex transmission risks posed by traditional burial and caregiving practices. Despite the availability of COMBI guidelines completed just prior to the Ebola outbreak, standard operating procedures developed in Sierra Leone for implementing integrated social mobilization interventions could not be readily instituted partly due to weak capacity to operationalize the guidelines and coordinate activities at a large-scale [7]. In the context of an already fragile health system, the Ebola outbreak undoubtedly introduced new and unique challenges that the country was ill-prepared to handle [8].

Early messaging overly emphasized Ebola as a ‘killer disease’ but fell short in providing actionable information on prevention, treatment, and possible survival [8]. Initial emphasis on fear, as well as lack of sensitivity to community values and traditions, contributed to people hiding from authorities and failing to seek medical care [9]. This reflected experiences from previous outbreaks in Africa [10]. At the same time, early anthropological research in Sierra Leone found that communities were willing to change behaviors and accept response measures such as safe burials if they were appropriately and continuously engaged [11, 12]. In August 2014, a national assessment of public knowledge, attitudes, and practices found that Ebola awareness and knowledge were already high in Sierra Leone; however, misconceptions, stigma, and other barriers were prevalent [9].

It is against this background that five partner organizations - GOAL, Restless Development Sierra Leone, FOCUS 1000, BBC Media Action, and the US Centers for Disease Control and Prevention (CDC) - developed an integrated, community-led, data-driven approach, with its core component consisting of large-scale community engagement in support of outbreak containment. The Social Mobilization Action Consortium (SMAC) was established in September 2014 and became operational in October 2014 in support of the Sierra Leone Ministry of Health&Sanitation’s Social Mobilization Pillar.

In this paper, we describe SMAC’s approach for community engagement that was implemented in Sierra Leone as part of outbreak response. We analyze and share over 50,000 semi-structured weekly reports from our network of Community Mobilizers. We draw upon this extensive data, and upon the collective experience implementing this integrated approach, to identify key lessons and make recommendations for future design, implementation and research.

## Taking Community Engagement at Scale in an Emergency

The SMAC program was directly implemented in approximately 70% of communities across Sierra Leone and covered all 14 districts, complemented by near universal radio and religious leader coverage. GOAL and Restless Development trained and supported nearly 2,500 community mobilizers who worked with communities to implement community action planning. Figure 1 shows the number of community visits per day. FOCUS 1000 trained and engaged 6,000 religious leaders from 2,000 mosques and churches to promote key messages and role model promoted behaviors, especially around safe burials. BBC Media Action supported 42 local radio station in 14 districts to improve the quality and synchronization of radio programming.

**Figure 1:**
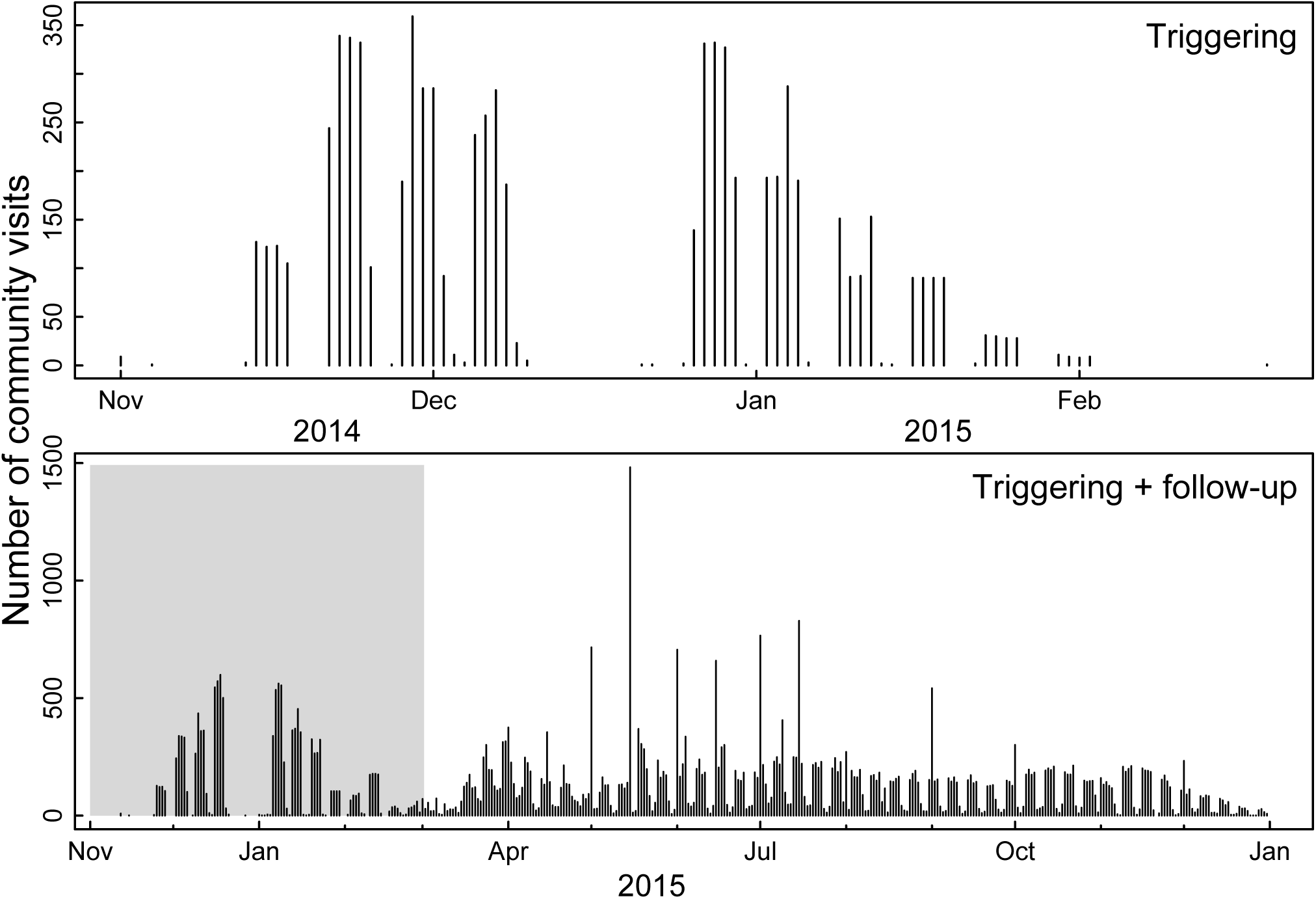
Community visits over time. Figure shows the number of community visits per day for the triggering events (top panel) and for the triggering and follow-up visits (bottom panel).

Community Mobilizers were recruited from an existing cohort of community health workers (CHWs), former Restless Development youth volunteers, and people nominated by their communities. Mobilizer community engagement was facilitated through the standardized Community-led Ebola Action (CLEA) approach (see the Table and Supporting Information) [13]. CLEA draws upon Participatory Learning and Action programming in HIV and AIDS contexts [14] and Community-Led Total Sanitation [15] within a structured community engagement approach based on Restless Development Sierra Leone’s Volunteer Peer Education Program.

CLEA departs from one-way health education and focuses on bottom-up community planning comprising triggering events and regular follow-ups (Supporting Information). The aim of CLEA triggering events was to create a sense of urgency, a desire to act, and local ownership. In each community, an initial triggering event facilitated by Mobilizers helped community members to undertake their own Ebola outbreak self-appraisal and analysis. Triggering resulted in the development of community action plans and identification of emergent Community Champions. These plans consisted of action points, often in the form of by-laws (such as restricting entrance to, and exit from, a community) that were implemented by communities and progress was monitored by community mobilizers.

Regular follow-up visits by Mobilizers supported maintenance of agreed actions within communities. Mobilizers captured both quantitative and qualitative data gathered from their engagements with communities during subsequent follow-ups using a standardized form (Supporting Information). Quantitative items included community surveillance metrics such as suspected numbers of Ebola cases reported, number of suspected cases alerted to authorities within 24 hours, number of survivors, number of suspected deaths, number of safe burials, and number of burials directly conducted by the community outside of the safe burial system. Qualitative items were captured through open-ended questions included commonly expressed concerns, Ebola risk perceptions, and narratives on community action plans.

Data were collected using paper-based forms across all districts from December 2014 through to September 2015, while a subset of the data from April to September 2015 were collected using a digital system in five active transmission districts (Western Area Urban, Western Area Rural, Port Loko, Kambia, Moyamba and Kono) using Open Data Kit (opendatakit.com). Starting in April 2015, digital data reporting incorporated religious groups and radio stations in the five aforementioned districts. However, in this paper, we only present data from Community Mobilizers utilizing the CLEA approach to allow for consistency in reporting format and method of data collection.

**Table 1:**
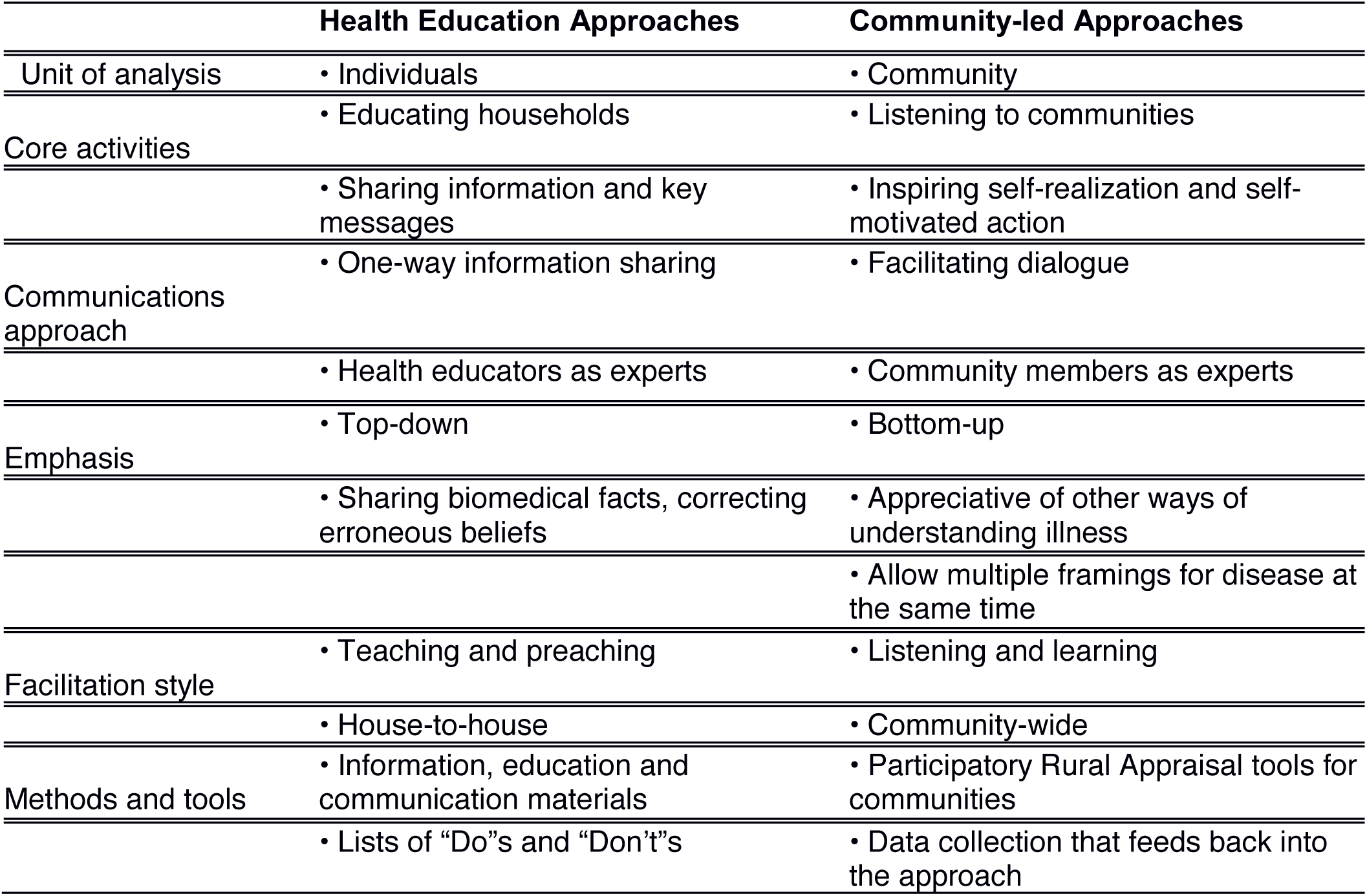
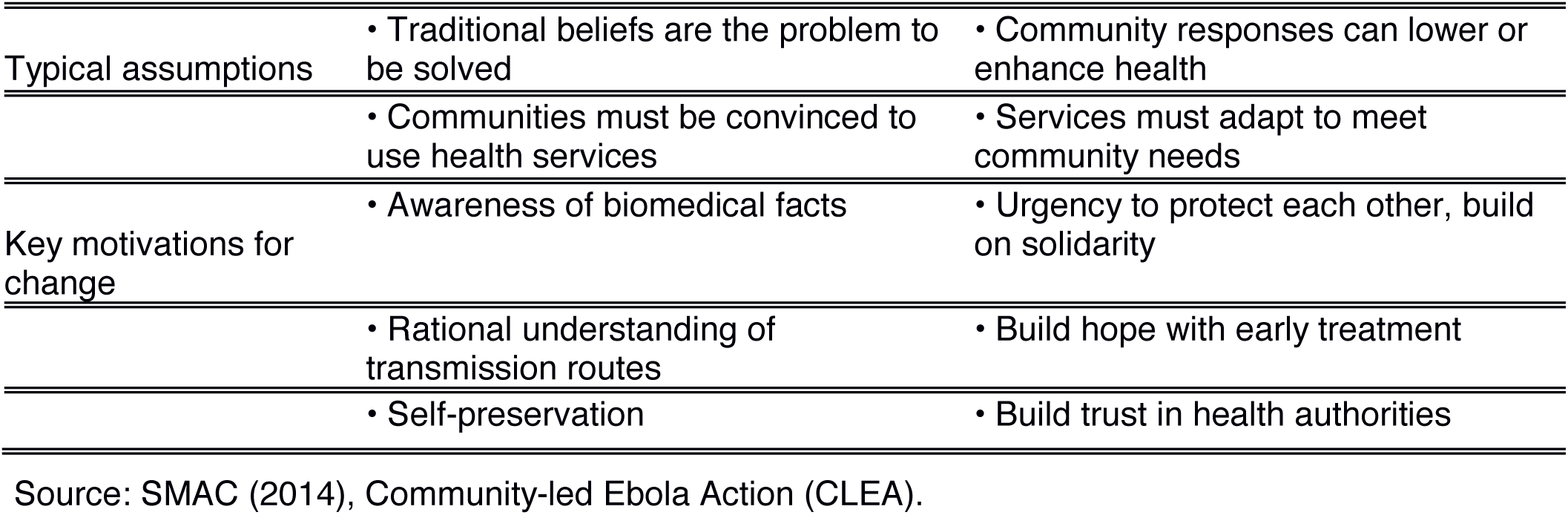
Comparison between Health Awareness and Community-led Approaches.

Using standardized monitoring forms, quantitative epidemiological data were collected on the total number of Ebola cases, number of cases referred to a health facility within 24 hours, number of survivors, number of suspected deaths, number of safe burials, number of burials conducted by the community, and the time elapsed since last suspected case. All quantities were compiled separately for males/females and children/adults. Knowledge, attitudes and practices of the communities were captured through a set of open questions, including but not limited to the following:

1. What are the most commonly expressed Ebola-related concerns expressed by community members?
2. What were the most commonly asked questions by community members?
3. What did the community initially assess and rank as key risks for contracting Ebola?
4. What bylaws have been developed on Ebola in this community?

The impact of community engagement can be quantified through health-related behaviours: referrals to health facilities, safe burials, and new cases identified. The other quantitative impacts are investigated in Fig. 2.

**Figure 2:**
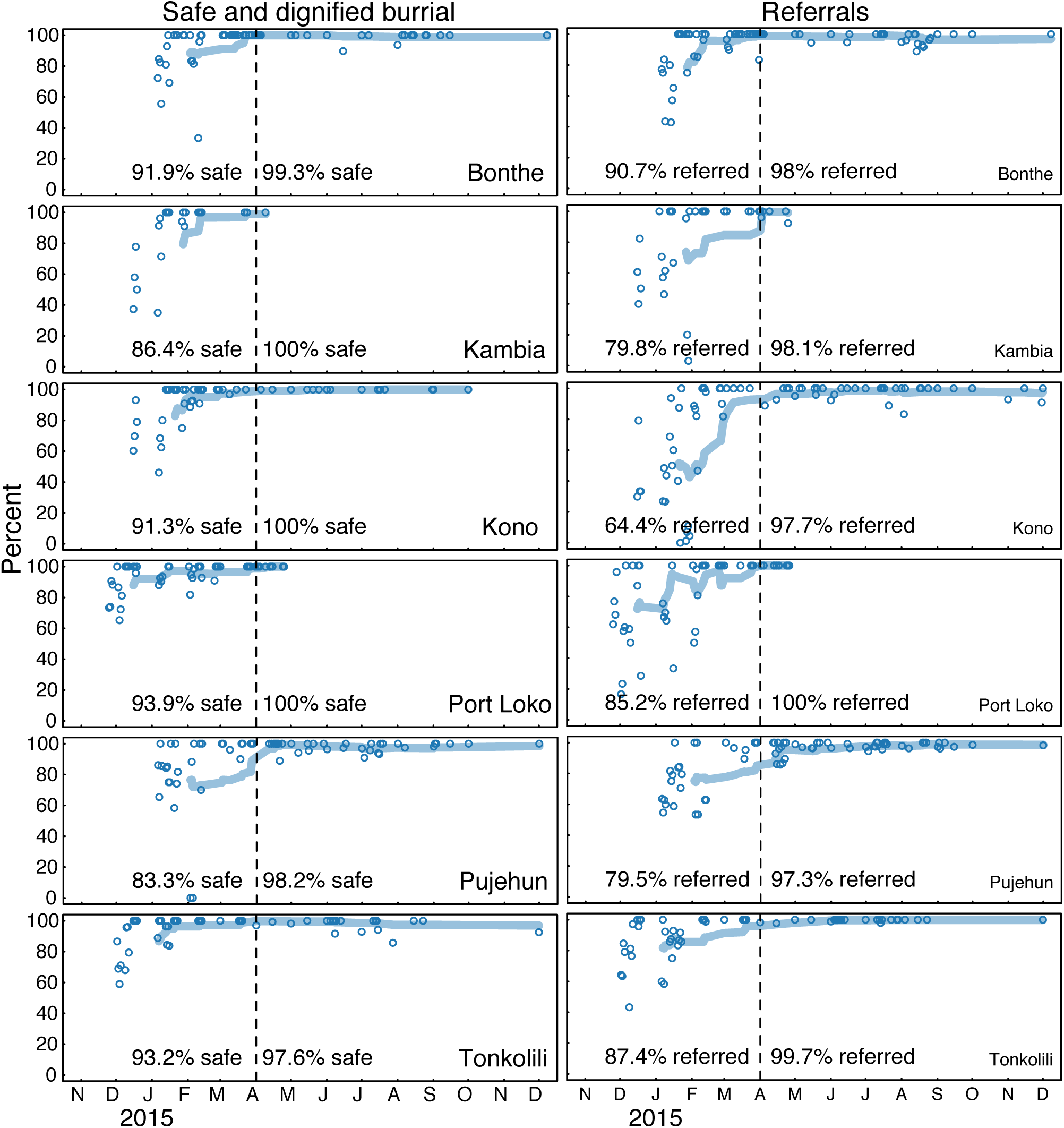
Behavioural impacts of community engagement. Increase in fraction of safe burials following deaths (right) and fraction of cases referred to a health facility with 24 hours (left). We divide the data per district and plot our estimates for percent of cases referred, and percent of safe burials following deaths, at different visits. The dotted line show the transition from period 1 (paper based) and period 2 (digital) which also after most community were already triggered and follow-ups ramped up.

## Insights gleaned from monitoring data

Through the CLEA approach, Mobilizers made multiple visits to >12,000 communities nationally. Community Mobilizers using the standardized paper forms engaged 2,113,902 community members over this time period, of which 50.2% were female while 49.8% were male, and 46% were young people while 54% were adults. These numbers do not represent the number of unique individuals, but consider all interactions as a result of repeat visits to communities. During triggering events, the average number of participants per community was 48; in follow-up visits however, the number more than doubled to 113 participants per visit. The demographics of participants did not significantly change from triggering events to follow-up visits. In parallel, using digital reports, Mobilizers visited households/neighborhoods individually and engaged with over 3,129,380 non-unique community members. Similar to the community visits, 52% of these were female and 48% male. The average visit consisted of an interaction with 57 community members with most around 25 but some as high as hundreds.

The fact that average number of community members engaged per community more than doubled between triggering events and follow-up visits suggest important buy-in and support from the communities. Moreover, monitoring data revealed that 100% of communities developed community action plans containing 3 or more action points on average. Between April and September 2015 when monitoring was fully operational including through the digital system, Mobilizers followed-up on 63,110 cumulative action points. Of these collective action points, 85% were assessed as “in progress” while 7% were marked as “achieved” and another 7% were “not achieved.” Some summary statistics from our data on action points and bylaws collected through the paper forms are presented in Fig. 3.

**Figure 3:**
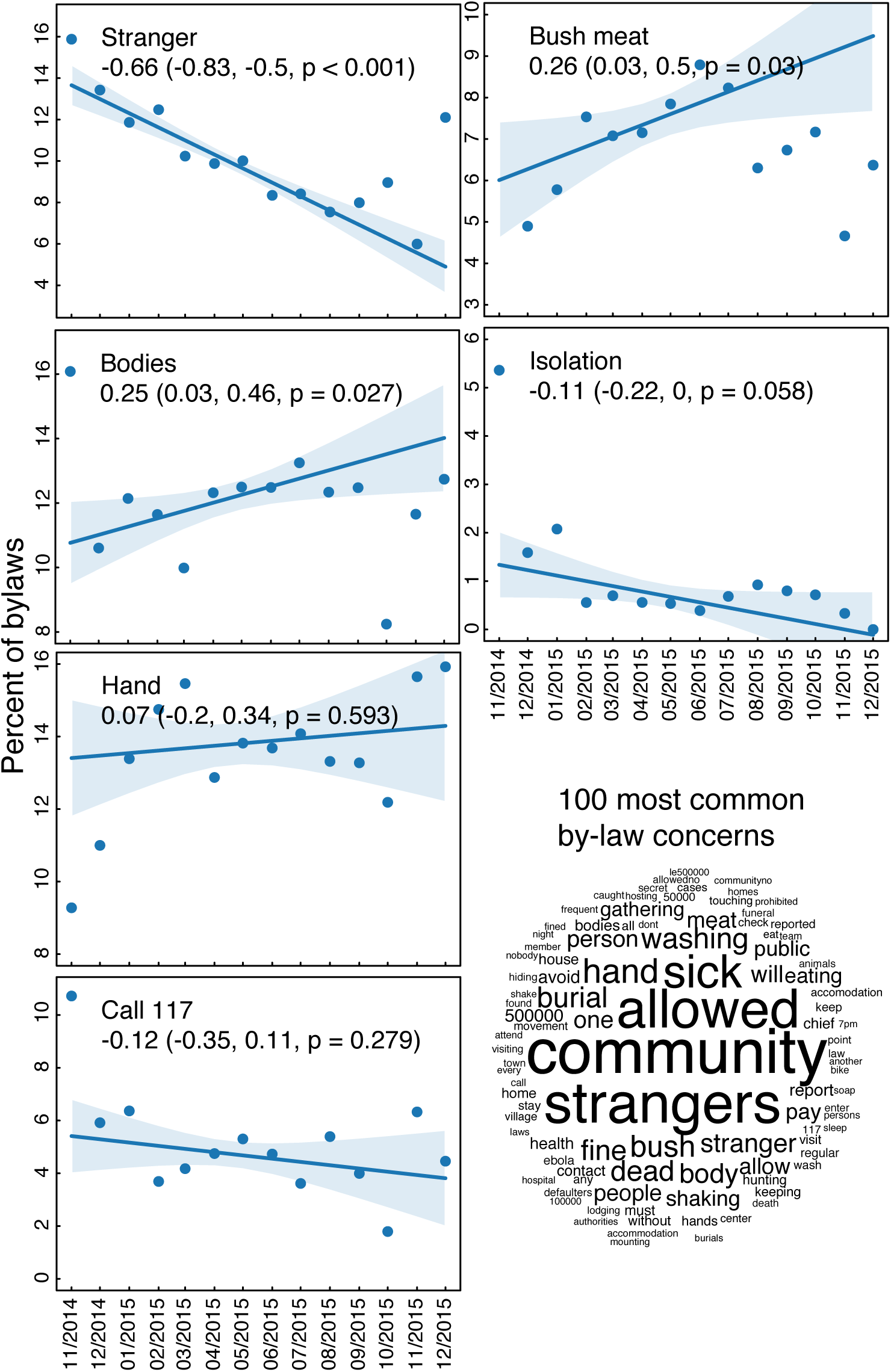
Content of community bylaws. (left) Frequency of common topics mentioned in community bylaws over time during follow-up visits with regressions weighted by the number of bylaws in the month. (bottom right) Qualitative representation of the most common concerns and topics in all community bylaws. Numbers refer to the toll-free national alert system (117) and to fines associated with the bylaws (e.g. Le 500,000 ≃$60 in United States dollar).

The main behavioral outcomes measure of the community action planning visits and follow-ups were (i) timely referrals of sick household members for medical care and (ii) timely requests of safe burials for deceased family members (Fig. 2). In our analysis, we divided the data by district and plotted our estimates for percent of cases referred, and percent of safe burials following deaths, at different visits. The results indicated an increase over time in fraction of safe burials following reported deaths and fraction of reported cases referred for medical care within 24 hours. The qualitative data were categorized and themes examined. We then calculated the frequency of common topics mentioned in community bylaws over time with regressions weighted by the number of bylaws in the month (Fig. 3).

The data show shifts in action points prioritized and implemented by community members during the intervention period (Fig. 3). For instance, by-laws around allowing isolation of communities and consumption of bush meat declined steadily and statistically significantly from November 2014 to December 2015, while by-laws around dead bodies or hand washing increased statistically significantly over this same period.

## Reflections on Lessons Learned

### Community Engagement, Real-time Data Collection and Action Planning at Scale

Results demonstrate the achievement of large-scale community engagement at national level in a health emergency context. They also demonstrate that community-level engagement based on a structured, participatory, and monitored methodology can support the collection of real-time data at scale, the establishment of meaningful feedback loops for both upward and downward exchange of information, along with collective planning and prioritization of actions at the household and community level. That this dataset was collected by Community Mobilizers and their communities echoes previous studies that have shown that digital data collection can be successfully implemented by community health workers with little experience if adequately trained and supervised [16].

### Impacts on Addressing High-Risk Practice

Results found a correlation between communities engaged with the CLEA approach and increasing trends in use of safe burials and early referral of sick people over the course of the outbreak in Sierra Leone. Early referral was a key action point within the community action plans in the triggered communities and communities we analyzed correlated with a significant increase in in 24-hour referrals (Fig. 2).

While it is important to note that the SMAC program coincided with a plateau and decrease of the epidemiological curve and overall increase in response resourcing - including a more responsive Ebola hotline (117), increased number and professionalism of burial teams and number of Ebola Treatment Centers - the results do demonstrate that community action plans included a range of community-specific controls and actions. Communities reported 90% of collective action points either achieved or in progress/maintained.

### Community-based Surveillance and the Integration of Community Engagement

The data show that Community Mobilizers and religious leaders became active agents in Ebola surveillance at national level. Although community surveillance was not initially integral to the original CLEA model, it soon became a core component driven by local needs and level of trust established between SMAC and the target communities. Mobilizers made an average of 133 community visits per day nationally using paper forms and 151 visits per day nationally using digital reports. More than 1,500 mobilizers received SIM cards and access to free mobile phone calls via a SMAC Closed User Group. All Mobilizers were trained in alerts mechanisms within District response authorities (i.e. reporting of potential cases or deaths). These factors essentially established a de facto community surveillance system and closed feedback loop that resulted in SMAC Community Mobilizers making more than 2,500 alerts to response agencies at district level [17].

Community engagement spans both demand generation and ensuring that the supply of essential services meets increased demand. Therefore, response actors should place more emphasis on creating strong functional linkages between community-level prevention and other aspects of the response, particularly surveillance efforts. The data affirm the belated recognition in the Standard Operating Procedures for the Social Mobilization Pillar and Surveillance Pillar (along with all other biomedical pillars) that closer integration is integral to an effective Ebola response [7].

### Integrated Communication

The SMAC model provides further evidence that behaviour change interventions are most likely to be effective when a combination of communication channels and platforms are appropriately used, combining community-based interpersonal communication with mass media, and working in support of government policies. This approach is more likely to achieve rapid behaviour change in an outbreak setting, as consistent information and messaging that support community-led responses are repeated and reinforced via multiple channels, thereby increasing information credibility and reducing confusion caused by mixed messaging.

Radio provided a platform for the participation of trusted messengers such as religious leaders, community champions, and traditional healers as well as survivors and responders to clarify messages and to discuss concerns.

Religious leaders trained and engaged through SMAC were able to leverage their wide-reaching network across all parts of the country to persuade communities to adopt behavioural changes especially modifications in traditional burial practices. Engagement of traditional healers – when the outbreak was waning but Ebola transmission continued to be linked to unsafe healing practices.

## Limitations

One of the main shortcomings of the SMAC dataset is that data from December 2014 through December 2015 were collected through paper reports, while a subset of data from April through September 2015 were collected directly through digital reports. Merging these datasets proved problematic

However, developing a digital data collection system proved invaluable in ensuring information on behaviours in communities was immediately available. Digital data collection overcomes the limitations of paper-based data collection, including collection, transportation, and onerous and error-prone data entry. That being the case, issues of charging devices, connectivity and mobile network reliability can hinder and frustrate digital data collection in setting such as Sierra Leone.

CLEA is a methodology that prioritizes community discussion and limited, targeted household engagement over blanket house-to-house (H2H) engagement. However, both SMAC operational and external factors required adaptation of the CLEA approach and explain significantly higher non-unique community member engagement in primarily Western Area and to some degree in Kambia. In Western Area, large compounds contained a cluster of households/families; and mobilizers at times supported ‘surge’ campaign activities directed by the Social Mobilization, aligning to Social Mobilization Pillar H2H methodologies and working in and around quarantined areas. Mobilizers were also integrated into health and contact tracing teams to provide support to geographically targeted H2H visits.

It should be noted that data was self-reported by communities and collected by community mobilizers which may have resulted in reporting bias.

## Looking to the Future

Data from the respective SMAC interventions reveal that it is feasible and cost-effective to support communities to plan for and monitor their own actions in a quantifiable way during an epidemic, provided the right enabling and reinforcing structures are in place. These include: strong baseline data identifying key behavioural determinants; regular and timely system for capturing and reporting monitoring data; systematic and consistent community engagement approaches emphasizing two-way communication and feedback loops; continuous supervision and ongoing peer-to-peer support for community audiences; and adequate logistical and communication support. Furthermore, the data suggests that communities are capable of engaging in localized surveillance and referral if given the right tools, support and linkages to the formal health structures/systems.

There are unique opportunities for future analyses of the large-scale data collected through the SMAC intervention. Potential areas for future analyses include: (i) Exploration of correlations between reported behavioural trends and changes in disease epidemiology. Coupling of the SMAC dataset with disease models could provide potential opportunities to ascertain correlations and test hypotheses about the behaviours most likely to impact Ebola spread. (ii) Comparison and triangulation of SMAC community surveillance data with data from other pillars, particularly surveillance and burials, to understand how community-reported data compares to data collected through other sources. (iii) Further coding and analysis of the qualitative weekly data collected by mobilizers, including details on rumors, reported changes, survivor acceptance, challenges and common concerns, and action points.

Overall, given its size and scope, the complete SMAC data provide several unique opportunities for multidisciplinary research to better understand and quantify the health-related effects of specific events and behavioural interventions.

