## Supplementary Information for "Community Engagement during outbreak response: standards, approaches, and lessons from the 2014-2016 Ebola outbreak in Sierra Leone"

**Acknowledgements**

The Social Mobilization Action Consortium (SMAC) partners include Centers for Disease Control, Focus 1000, BBC Media Action, GOAL and Restless Development. SMAC worked within the strategy of the Ministry of Health and Sanitation’s National Social Mobilization Pillar and is funded primarily by United Kingdom Department for International Development and the Bill and Melinda Gates Foundation. A proportion of Restless Development community mobilisers were also funded by Age International and UNICEF. BMA also thanks Bill and Melinda Gates for their support of this work and their sponsorship through the Global Good Fund. LHD acknowledges support from the National Institutes of Health and 1P20 GM125498-01 Centers of Biomedical Research Excellence Award.

**SMAC’s Integrated Community Engagement Platform.**

In August 2014, a national assessment of public knowledge, attitudes, and practices (KAP-1) found that initial awareness raising and knowledge creation efforts had been successful, however, misconceptions, stigma and other barriers continued to prevent good behaviors (Jalloh, et al.).Based on these findings, the Social Mobilization Pillar (SM Pillar) with support from FOCUS 1000, US Centers for Disease Control and Prevention and other partners developed a National Communications Strategy for Ebola Response in Sierra Leone (National Social Mobilization Pillar, 2014) as well as a comprehensive messaging guide (National Social Mobilization Pillar, 2014).

As the outbreak accelerated, there was growing recognition that community engagement should be a fundamental element of the response, serving as an essential demand driver for key services, such as medical care and safe burials, whilst providing feedback loops to help improve these services based on community needs and perceptions (Pedi, et al.). It is against this background that five partners from the SM Pillar – GOAL, Restless Development, FOCUS 1000, BBC Media Action, and CDC – developed a community engagement intervention to provide a coordinated, integrated and national approach to community engagement, whilst supporting the Sierra Leone Ministry of Health and Sanitation’s National Social Mobilization Pillar at national and district levels. SMAC was established in September 2015 with endorsement from MoHS, and initial funding from the United Kingdom Department for International Development.

SMAC’s Theory of Change was premised on the belief that communities are not passive recipients of health messages and services: They can and should be seen as equal partners in an emergency response. Communities are capable of assessing their risks and identifying mitigation steps through collective action planning, and willing to modify deeply-rooted socio-cultural practices and adopt new practices if empowered to do this on their own terms. To ensure this was based on two-way communication through trusted, meaningful and reliable mechanisms, SMAC prioritized an integrated, multi-platform approach, where multiple and reinforcing sources of communication and feedback were designed and implemented.

Key elements included:

- Provision of technical and secretariat services to the National Social Mobilization Pillar and District Social Mobilization Committees;
- Establishment and coordination of an integrated package of community engagement and social mobilization activities including:
  - 2,500 Community Mobilizers implementing CLEA;
  - Support to 42 Radio Stations;
  - Training and support for 2,000 Religious Leaders.
  - Engagement and support of Ebola survivors.
- A coordinating secretariat that worked to ensure integration of all elements of thew SMAC consortium, alongside with the Social Mobilization Pillar and other response pillars guiding the biomedical response.
- Real-time data collection and analysis of all interactions undertaken by community mobilizers, complemented by real-time data collection by religious leaders and radio stations.
- Development of weekly national and district level Situation Reports for social mobilization and community engagement which included: i) key challenges; ii) community observations; iii) district level actions/observations; iv) number of triggering and follow-up visits to communities; v) national and district social mobilization pillar updates; vi) death and sick alerts made by community mobilizers.

**Community-led Ebola Action (CLEA) Mobilizers**

Mobilizer community engagement was facilitated through the Community-led Ebola Action (CLEA) approach (Social Mobilization Action Consortium, 2014), designed and implemented by SMAC partners in consultation with staff and volunteers.

The CLEA approach involved assigning mobilizer pairs to a specific number of communities. In each community, an initial Triggering Event facilitated by mobilizers helped community members to undertake their own EVD outbreak self-appraisal and analysis. When successful, CLEA triggering created a sense of urgency and a desire to act. Triggering resulted in the development of community action plans and identification of emergent Community Champions. After triggering, regular follow-up visits by mobilizers supported maintenance of agreed actions within communities. Follow-up sessions included a combination of community discussions and targeted household visits. CLEA marked a departure from health education approaches.

The CLEA methodology prioritizes community discussion and limited, targeted household engagement over blanket house-to-house (H2H) engagement. However, both SMAC operational and external factors required adaptation of the CLEA approach and explain high household visitation found primarily in Western Area and to some degree in Kambia. In an urban context, adaptations to the CLEA methodology were made due to lack of community cohesion, high density informal urban settlement conditions and high mobility. For example, while Sierra Leone has an overall population density of 79/sq km, while Western Area, comprising the capital city Freetown and surrounding suburbs, has a population density of 1,224/sq km. Large compounds contained a cluster of households/families; and mobilizers at times supported ‘surge’ campaign activities directed by the Social Mobilization Pillar, aligning to Social Mobilization Pillar H2H methodologies and working in and around quarantined areas.

Adaptations used in this region included finding alternative champions/influencers in the absence of more traditional leadership structures; developing methodologies for demarcating communities; and using dialogue through household visits.

Key elements included:

- Prioritization of recruitment of CHWs, young people with volunteer experience, community members nominated by communities themselves;
- Comprehensive training on the CLEA methodology, interpersonal communication, Ebola transmission, alerts mechanisms, safety and security and data collection;
- Assignment of mobilizer pairs (one male and one female) to between 6 – 8 communities, to be triggered and follow up in rotation;
- Consultation with District authorities for identification of priority communities and with communities themselves to seek permission for entry and initiating engagement.
- Payment of stipend and reimbursement of travel expenses for all mobilizers, subject to work agreements and codes of conduct signed by mobilizers and implementing agencies;
- Weekly meetings of mobilizers to submit data, receive top-up training and report feedback;
- Submission of monitoring forms signed off by community leaders, including contact details of community representative for verification purposes;
- Provision of SIM cards and free phone calls via a Closed User Group for mobilizers.

**Religious Groups**

A high proportion of religious leaders continuously promoted key Ebola messages during Friday and Sunday prayers, while their mosques/churches also provided a range of support to quarantined families and undertook other community outreach events. There is a mosque and/or church in every community in Sierra Leone, and over 95% of Sierra Leoneans identify as a Muslim or Christian (Statistics Sierra Leone, 2004). SMAC recognized the need for religious leaders to role model safe practices and to provide social support to their peers and congregants, especially in shifting from traditional to Safe and Dignified Medical Burials (SDMB). FOCUS 1000 revitalized the Islamic Action Group (ISLAG) and the Christian Action Group (CHRISTAG) platforms as trusted channels of health communication, formerly leveraged in the late 1980s to achieve universal childhood immunization (UNICEF, 1996).

A national interfaith taskforce of 25 each of Islamic and Christian Action Group members was established in collaboration with the Sierra Leone Inter-Religious Council. The taskforce identified texts from the Quran and Bible to support the key messages on safe burials, early treatment, and acceptance of survivors and health workers. A total of 1,989 imams and pastors were trained, as well as 4,000 women and youth leaders from 2,000 mosques and churches across the 14 districts in Sierra Leone. A total of 28 faith-based district coordinators – one each of Islamic and Christian Action Group members in each district – were appointed. Similarly, 298 chiefdom coordinators and over 1500 section coordinators were appointed, reaching every section in the country. Such coordinated structure provided a trusted national network for engaging faith-based communities. The trained religious groups – under the leadership of appointed imams/pastors – role modeled preventive behaviors, promoted key practices, and served as trusted messengers in more than 300 radio and television programs.

*Activities and reach of religious leaders (April – September 2015)*

A total of 4,402 weekly reports were submitted by the targeted religious groups (mosques/churches) in the five districts using the digital data collection system. Of these, 53% were from mosques and 47% from churches. Data revealed that 60% of religious leaders promoted SDMB during sermons, 57% promoted early treatment, and 62% promoted acceptance of survivors and health care workers. Nearly 4 in 10 religious leaders per week participated in SDMBs. Religious leader participation in SDMB varied significantly across districts. It was higher in Kono (45%) and Port Loko (44%) than in Kambia (38%), Moyamba (33%), and Western Area (29%). On average, religious leaders participated in 2 SDMB per week with no significant differences across districts. In addition, 29% of mosques/churches provided support to quarantined homes, whereas 32% carried out other EVD community outreach.

**Radio**

The KAP-1 survey found that radio had by far the widest reach when compared to all other single channels of EVD information, with 88% of respondents reportedly receiving information on EVD through radio. More precisely, 79% respondents reported listening to the radio daily, while 13% listened weekly. BBC Media Action supported 36 Partner Radio Stations to produce and broadcast approximately 40 hours of EVD programming per week reaching all 14 districts. Partner Radio Stations produced and broadcast local radio programs, providing district-specific information in local languages to audiences. In addition, BBC Media Action produced two national radio discussion programs and a radio drama. Kick Ebola Live was a two-hour weekly magazine program with interactive SMS component, which was picked up by over 30 radio stations across the country via simulcast. Kick Ebola Nar Salone was a 30 minute pre-recorded show produced weekly and distributed to 36 partner stations across the country. And the drama entitled “Mr. Plan Plan” encouraged families to face dilemmas, discuss options and develop a plan.

BBC Media Action’s partner radio stations collaborated with other SMAC partners to ensure that trusted figures from the community, including representatives from the DERC, religious leaders, mobilizers and Community Champions, participated in radio discussion programs, providing up-to-date and accurate information and a range of different perspectives.

*Activities and reach of partner radio stations (April – September 2015)*

SMAC’s partner radio stations submitted 120 weekly reports. They collectively produced 189 radio programs (mean of 2 weekly) across the five districts on multiple EVD-related issues, including prevention, treatment, and transmission. Nearly all radio programs (99.5%) featured a community leader as a live guest or through recorded segments. Collectively, the programs generated 1,619 text messages from listeners. The majority of listeners’ text messages (70%) were related to locally driven issues on the effectiveness of the response, the impact of EVD on education, youth welfare, and employment. In addition, 25% of messages were related to survivors, 22% on burials and 20% on re-infection. Given the small sample size for radio programs, district-level analysis is not appropriate.

SMAC’s partner radio stations provided a channel for reaching large proportion of the population with key EVD information. Radio provided a platform for the participation of trusted messengers such as religious leaders, community champions, and traditional healers as well as survivors and responders to clarify messages and to discuss concerns. Listeners were able to send text messages or call-in to ask questions and share their views. In the earlier stages of the response, rumors and misconceptions were often amplified via low quality radio programming. SMAC supported 36 national and local radio stations through training and mentoring to strengthen the quality of weekly programs in local languages. By improving the quality of content, SMAC was able to engage audiences and contribute to the accuracy of information disseminated via radio. Using trusted sources on radio, the messages were more likely to resonate with the local populations.

This was coupled with 189 radio programs^[1]^ wherein community leaders were almost always featured as guests.

**Supplementary Table 1. Percent distribution of religious leader activities, and text messages from radio programming, and concerns expressed by communities during mobilizer visits, April – September 2015**

| **Indicator** | **Western Area** | **Kambia** | **Port Loko** | **Kono** | **Moyamba** | Total |
| --- | --- | --- | --- | --- | --- | --- |
| **Weekly promotion of key EVD messages by SMAC trained religious leaders (N=4,402)** | | | | | | |
| Safe and dignified medical burials | 63.4% | 62.4% | 62.8% | 53.8% | 51.5% | 59.5% |
| Early medical treatment | 62.1% | 63.0% | 60.7% | 51.6% | 41.6% | 57.0% |
| Respect for Ebola survivors and health care workers | 71.7% | 59.9% | 66.7% | 54.0% | 51.6% | 62.2% |
| **Outreach activities by trained religious groups (N=4,402)** | | | | | | |
| Provided support to quarantined homes / families the community | 28.5% | 31.7% | 47.2% | 15.7% | 2.8% | 28.8% |
| Carried out other community outreach on Ebola | 34.6% | 28.4% | 39.0% | 32.4% | 18.5% | 32.4% |
| **Content of radio programs (N=120)** | | | | | | |
| Safe and dignified medical burials | 22.9% | 0.0% | 15.8% | 25.0% | 17.9% | 19.2% |
| Stay safe while you wait | 20.8% | 16.7% | 5.3% | 12.5% | 5.1% | 12.5% |
| Survivors | 27.1% | 0.0% | 10.5% | 0.0% | 7.7% | 15.0% |
| Stigmatization/discrimination | 27.1% | 0.0% | 5.3% | 0.0% | 2.6% | 12.5% |
| Re-infection | 8.3% | 0.0% | 10.5% | 0.0% | 15.4% | 10.0% |
| Other | 64.6% | 83.3% | 73.7% | 87.5% | 82.1% | 74.2% |
| **Content of text messages received during radio programs (N=120)** | | | | | | |
| Safe and dignified medical burials | 25.0% | 0.0% | 10.5% | 25.0% | 25.6% | 21.7% |
| Stay safe while you wait | 25.0% | 16.7% | 10.5% | 25.0% | 5.1% | 15.8% |
| Survivors | 37.5% | 0.0% | 10.5% | 12.5% | 23.1% | 25.0% |
| Stigmatization/discrimination | 37.5% | 0.0% | 5.3% | 0.0% | 7.7% | 18.3% |
| Re-infection | 29.2% | 0.0% | 10.5% | 12.5% | 17.9% | 20.0% |
| Misappropriation of Ebola funds | 4.2% | 0.0% | 0.0% | 0.0% | 2.6% | 2.5% |
| Other | 56.3% | 83.3% | 84.2% | 75.0% | 76.9% | 70.0% |

1. Limited to the 120 samples submitted using the Digital Data Collection System between April and September 2015, and only in the 5 districts. A substantially higher number of radio programs were produced with support from SMAC partner BBC Media Action across the 14 districts and over a longer period.
